# Bulb bankruptcy: energy dynamics in *Eucomis autumnalis*, a South African mesic grassland geophyte

**DOI:** 10.64898/2026.07.02.735992

**Authors:** Craig D Morris, Sindiso Nkuna

**Affiliations:** Agricultural Research Council – Animal Production (ARC-AP), c/o School of Agriculture and Science, University of KwaZulu-Natal, Private Bag X01, Scottville, 3201, South Africa; School of Agriculture and Science, University of KwaZulu-Natal, Private Bag X01, Scottville, 3201, South Africa

**Keywords:** autumn, carbohydrate reserves, defoliation, geophytic forbs, underground storage organs

## Abstract

Intense grazing and trampling by livestock reduce forb populations in South African mesic grasslands by limiting photosynthesis, thereby depleting carbohydrate reserves in underground storage organs (USOs) required for persistence. The effects of simulated herbivory on above- and below-ground growth, bulb starch reserves, and water content in the bulbous geophyte, *Eucomis autumnalis*, were assessed over a 468-day pot experiment, with seasonal dynamics described using Generalised Additive Models. Four intense summer and autumn defoliations reduced seasonally integrated above-ground dry matter (−83.7%), bulb mass (−51.7%), starch concentration (−32.3%), total starch pool (−68.9%), and bulb water pool (−35.1%), while increasing bulb water content (+29.4%) relative to controls. Bulb growth and starch pools (bulb mass × starch concentration) were most strongly suppressed during late summer and autumn, when control plants increased bulb mass and replenished starch reserves. Repeated defoliation forced plants to divert assimilates to regrowth, disrupting seasonal source–sink dynamics and driving progressive depletion of reserves; bulbs were consequently pushed towards energetic bankruptcy, potentially unable to sustain future growth and persistence under chronic grazing. These findings highlight the need to avoid high stocking densities, protect plants from intense autumn grazing, and provide periodic full-year recovery rests to conserve geophytic forbs in mesic grasslands.

## INTRODUCTION

Grasslands are among the most hidden ecosystems, with between 60 to over 95 per cent of their biomass residing underground as roots, rhizomes, and other plant organs (Stanton 1988; Mokany et al. 2005). High root-to-shoot ratios (RSR) are characteristic of seasonal and disturbance-prone biomes, where recurrent fire, frost, and drought favour allocation to belowground structures (Jackson et al. 1996; Qi et al. 2019; Ottaviani et al. 2020). In South Africa, for example, recent estimates for subtropical grasslands report RSR values far exceeding the global average of 3.22, particularly in fire-dependent mesic grasslands, where the RSR approaches 20 (Slooten et al. 2026). This highly skewed allocation in mesic grasslands (>650 mm MAP) translates into more than 50 t ha⁻¹ of dry plant material concentrated in the top 20 cm of soil, compared with just over 3 t ha⁻¹ above ground (Slooten et al. 2026).

Grasses exploit the soil for nutrients, water, and anchorage (Weaver 1958), but many herbaceous non-grasses (monocotyledonous and dicotyledonous forbs) in mesic grasslands of southern Africa have especially invested in life underground (Carbutt et al. 2017), producing roots as well as specialised structures and organs for storing energy, nutrients, and water, and enabling clonal expansion (Klimešová et al. 2018; Siebert et al. 2024). Such geophytes produce a range of perennial underground storage organs (USOs) metamorphosed from stems and roots (Klimešová 2025), including tubers, rhizomes, bulbs, corms, and lignotubers (de Moraes et al. 2016; Orzell et al. 2024; Martínková et al. 2025). These USOs enable plants to survive seasonal frost, fire, and periods of water and nutrient limitation, as well as the occasional loss of aerial parts due to unpredictable disturbances such as herbivory (Fidelis et al. 2014; Lubbe et al. 2021). Channeling carbon acquired through photosynthesis, together with water and nutrients from the soil, into USOs represents an evolved resource allocation strategy that ultimately pays off, enabling geophytic forbs to persist in the long term in seasonally disturbed grasslands (Veselý et al. 2012; Tribble et al. 2021).

Cyclical seasonal processes of resource acquisition, storage, and production are crucial for maintaining and expanding USOs, thereby ensuring the longevity of forbs in grasslands. These metabolic processes are akin to maintaining a solvent business, following the useful economic metaphor proposed by Bloom et al. (1985), Bloom (1986), and Chapin et al. (1990) to describe plant storage and growth dynamics. In this framework, photosynthetic tissues function like productive assets, acquiring resources that can be reinvested into future growth. During favourable growing seasons, geophytes invest in leaves and roots while storing surplus carbon and nutrients in specialised tissues in USOs, much like profits saved as capital reserves. Although storage carries costs, these reserves act as both savings and insurance, sustaining plants through dormancy and financing reproduction and rapid spring regrowth (Morris 2021), thereby supporting long-term persistence in grasslands.

Herbivory during the growing season disrupts this resource economy by damaging the plant’s productive infrastructure (Morris 2021; Morris and Nkuna 2025a) and altering allocation priorities (Bloom et al. 1985; Chapin et al. 1990). Defoliation reduces photosynthetic leaf area, immediately lowering carbon income and limiting resources for growth, storage, and reproduction (Thomas et al. 2017). Herbivory can trigger induced resource sequestration — a tolerance response in which carbon and nutrients are rapidly shifted into storage organs to enhance regrowth — but repeated or intense defoliation may overwhelm this mechanism, leading to exhaustion of bulb reserves and compromised plant vigour and persistence (Orians et al. 2011). For grassland geophytes, repeated leaf loss through grazing or trampling (Chamane et al. 2017a) is particularly costly because summer is the main period for carbon sequestration (Morris and Nkuna 2025b). In economic terms, such herbivory reduces income while increasing compensatory expenditure on regrowth (McNaughton 1983). Plants draw on stored reserves to replace lost leaves and restore photosynthetic capacity, but repeated disturbance can progressively divert resources away from reserve replenishment towards short-term survival (Iwasa and Kubo 1997). This can ultimately lead to “bulb bankruptcy”, where reserves are depleted when replacement costs exceed seasonal income, reducing growth, vigour, resilience, and, ultimately, the persistence of grassland forbs in chronically overgrazed mesic grasslands (Scott-Shaw and Morris 2015).

We examined a key dynamic in the bulb bankruptcy framework: that repeated summer defoliation, simulating herbivory, disrupts seasonal carbohydrate cycling required for spring growth while sustaining below-ground reserves for regrowth in geophytic forbs. In a pot experiment, we tracked changes through two winters and one summer growing season in (1) foliar production, (2) bulb mass, (3) bulb starch concentration, and (4) bulb water content in the mesic grassland geophyte *Eucomis autumnalis*, comparing control and repeatedly defoliated plants. We also quantified total bulb pools (concentration × bulb mass), because reserve function depends on both concentration and storage organ size (Chapin et al. 1990; Janeček et al. 2015; Klimešová et al. 2017). We predicted that defoliation during the critical late-summer to autumn replenishment window would most strongly reduce reserve accumulation, thereby limiting overwintering capacity and subsequent spring growth, as proposed by Morris and Nkuna (2025b). The findings of this study can inform grazing management by highlighting critical seasonal windows when defoliation has disproportionate effects, supporting more adaptive livestock strategies aimed at sustaining below-ground reserves and maintaining diverse forb communities in mesic grasslands

## MATERIALS AND METHODS

### Study species

*Eucomis autumnalis* (pineapple lily), a member of the Hyacinthaceae, is a perennial deciduous geophyte occurring in wetter areas of mesic grassland in eastern and south-eastern South Africa. The species develops a large ovoid bulb (8–10 cm in diameter) and a basal rosette of broad, fleshy, soft-textured leaves with wavy margins, reaching up to 50 cm in length and 13 cm in width. Inflorescences are typically 50–60 cm tall (Pooley 1998; PlantZAfrica n.d.), and leaves senesce in winter, often removed by fire.

*Eucomis autumnalis* is widely utilised in traditional medicine, with extracts derived from bulbs, roots, stems, and leaves reported to treat a broad range of conditions, including pain, fever, urinary and respiratory ailments, inflammation, wounds, fractures, and teething (Taylor and van Staden 2001; Oyedemi et al. 2025; Ndlovu and Fisher 2026). Pharmacological studies further indicate potential applications in bone and cartilage regeneration in osteoarthritis (Alaribe et al. 2018). There are no documented reports of leaf grazing in *E. autumnalis*, although its upright growth form and soft leaves likely make it susceptible to trampling, particularly under dense livestock stocking (Chamane et al. 2017b).

### Study site and design

The pot trial was conducted under light hail netting (approximately 5% shading) at the N.M. Tainton Arboretum on the Pietermaritzburg campus of the University of KwaZulu-Natal, South Africa. The site is in a warm, humid subtropical climate with a mean annual rainfall of approximately 844 mm, summer maximum temperatures frequently exceeding 30–35 °C, and winter minimum temperatures around 3 °C (Morris 2021; Morris and Nkuna 2025a, 2025b).

Mature *E*. *autumnalis* plants, obtained from a commercial nursery, were grown in 8-litre pots filled with a nutrient-rich organic growing medium characterised by high organic carbon (>6%), low nitrogen (0.55%), slightly acidic pH (5.53), and elevated macronutrient concentrations (P = 81 mg L⁻¹, K = 156 mg L⁻¹, Ca = 2960 mg L⁻¹, Mg = 410 mg L⁻¹) (Manson et al. 2020). Watering was applied to ensure establishment and growth, including irrigation to saturation at the end of winter to trigger spring growth, with additional supplementary watering during periods of low rainfall and high temperature stress in summer to maintain growth.

The experiment followed a completely randomised design with two treatments comparing defoliated plants with undefoliated controls over two winters and one summer, from 20 June 2024 to 1 October 2025. Defoliation was imposed by cutting plants to 10 cm above the soil surface, thereby removing most leaf material, on four occasions during summer and autumn: 13 November 2024, 12 December 2024, 7 February 2025, and 10 April 2025. The first two defoliations were applied four weeks apart during peak summer growth, while the third and fourth were applied after longer intervals of about eight and nine weeks as growth rates declined later in the season. This treatment was intended to simulate the repeated leaf loss and mechanical damage imposed on herbaceous forbs by livestock trampling under high-density stocking in mesic grassland (Chamane et al. 2017a, b), and to assess how defoliation at different stages of the growing season influenced above- and below-ground growth patterns.

A total of 30 plants were grown and harvested destructively over the study period to quantify seasonal changes in bulb variables before and after defoliation commenced. Before treatment application in early summer, all plants followed a common growth trajectory and were sampled (n = 2 per date) for below-ground growth on three occasions: 20 June 2024, 14 August 2024, and 13 November 2024. Following the first defoliation on 13 November 2024, control and defoliated plants were destructively harvested (n = 2 per date per treatment) on six occasions: 12 December 2024, 7 February 2025, 10 April 2025, 17 June 2025, 31 July 2025, and 1 October 2025. All plants assigned to the defoliation treatment received all scheduled defoliations before harvest.

### Vegetation measurements

Above-ground dry matter yield (AGDM) was measured from aerial biomass (excluding occasional inflorescences), harvested from mid-November 2024 (when sufficient material was available) until the end of winter 2025, when all above-ground growth had senesced. At each harvest, aerial material was separated from bulbs and roots and dried at 60°C for seven days. Roots were removed from bulbs, which were cleaned and microwaved for 10 s at 800 W to deactivate starch-degrading enzymes (Chlumská et al. 2014; Morris and Nkuna 2025b), before determining bulb fresh mass. Bulb dry mass was measured after oven-drying at 60°C for seven days. Bulb water content was calculated as the difference between fresh and dry mass, expressed as a percentage of fresh mass. Starch content of dried bulbs, a key non-structural carbohydrate stored in the USOs of geophytes (Martínez-Vilalta et al. 2016), was determined using an enzymatic assay at the Plant Analysis Laboratory, KwaZulu-Natal Department of Agriculture, Cedara, South Africa (see Morris and Nkuna 2025b for details). Total starch and water pools were calculated by multiplying their respective proportions by bulb dry mass (Chapin et al. 1990).

### Statistical analysis

A regression-based approach was used to describe seasonal changes over time, with two replicate plants per treatment harvested at each sampling period and their mean values used for model fitting. Given the limited number of plants available, sampling was distributed as widely as possible across the season to capture expected fluctuations in growth dynamics, rather than increasing replication at fewer time points (e.g., Morris and Nkuna 2025b). The fitted models were used to identify peaks, troughs and seasonal patterns of change, and to compare derived variables (rates of change) and integrated seasonal responses (area under the curve) between treatments (Hunt 1979).

Generalised additive models (GAMs) were used to describe seasonal changes in all measured attributes. Although simple polynomial functions can also capture peaks and troughs, they often produce unrealistic curvature at the beginning and end of the fitted curves to maintain symmetry. GAMs instead use flexible smoothing functions to fit fluctuations without the constraints imposed by fixed polynomial parameters (Hastie and Tibshirani 1987), allowing complex seasonal patterns and non-linear trends to be modelled more realistically.

All statistical analyses and visualisations were performed in Python 3. Non-linear relationships were modelled with GAMs fitted as penalised cubic smoothing splines (k = 3), allowing underlying trends to be captured while limiting overfitting. The penalised smoothing splines used in this study were implemented through scipy.interpolate.UnivariateSpline, which is part of the Python SciPy library (Dierckx 1995; Virtanen et al. 2020). All six AGDM observations were used to fit separate curves for control and defoliated plants. For bulb variables, a common curve was fitted to the first three shared observations, after which separate curves were fitted for control and defoliated treatments using the six subsequent observations per treatment following the onset of defoliation. Ninety-five percent confidence intervals were estimated for all fitted curves using parametric bootstrap resampling (2,000 iterations). The Python code used to implement the GAMs was developed using Anthropic’s Claude (Sonnet 4.6), which provides a natural-language interface for building, running, and refining Python workflows and libraries. All graphical outputs were checked against raw data plots to ensure that fitted trends were biologically reasonable and consistent with the observed data.

Given the small sample size (n = 6), p-values associated with both the linear and smoothed components were not considered reliable for formal inference and are therefore not reported. Instead, model outputs are interpreted descriptively using the proportion of deviance explained (R²_dev_) as a relative goodness-of-fit metric, root mean square error (RMSE) as an estimate of prediction error, and, primarily, the visualised fitted curves with their 95% confidence intervals to show variation around the fitted curves, with emphasis on the biological plausibility of the modelled seasonal changes.

To calculate instantaneous rates of change (units per day) across the sampling period, particularly during key phases such as autumn when bulb reserves are typically replenished, first derivatives were obtained analytically from fitted GAM splines by refitting models with fourth-order smoothing splines, yielding smooth cubic derivative functions. Area under the curve (AUC) for each GAM was calculated by numerical integration over the full observation period. The AUC (unit·day or %·day) represents the total time-integrated value of each variable (AGDM, starch content, starch pool, water content, and water pool) across the season, summarising both its magnitude and duration in a single seasonal metric.

Relative effects of defoliation were expressed as percentage differences between defoliated and control plants. Effect sizes were calculated using Cohen’s *h* for proportional variables (starch content %, water content %) and Cohen’s *d* for continuous variables (mass and pool sizes), following Goulet-Pelletier and Cousineau (2018). Comparisons were based on paired post-treatment observations (n = 6 per treatment).

## RESULTS

The GAMs explained good to excellent proportions of the deviance (R²_dev_) in the seasonal dynamics for control plants across all variables: AGDM (R²_dev_ = 0.734, RMSE = 22.39 g), bulb biomass (R²_dev_ = 0.890, RMSE = 16.7 g), starch concentration (R²_dev_ = 0.794, RMSE = 3.83%), starch pool (R²_dev_ = 0.822, RMSE = 11.4 g), bulb water concentration (R²_dev_ = 0.918, RMSE = 2.11%), and bulb water pool (R²_dev_ = 0.678, RMSE = 7.13 g).

Model fits for defoliated plants ranged from excellent to weak: AGDM (R²_dev_ = 0.820, RMSE = 2.45 g), bulb biomass (R²_dev_ = 0.774, RMSE = 8.08 g), starch concentration (R²_dev_ = 0.512, RMSE = 8.61%), starch pool (R²_dev_ = 0.600, RMSE = 7.15 g), water concentration (R²_dev_ = 0.832, RMSE = 3.24%), and bulb water pool (R²_dev_ = 0.326, RMSE = 5.00 g).

Modelled seasonal patterns of aerial growth and each below-ground variable in the bulbs of *E*. *autumnalis* are described below for both control and defoliated plants, including how each variable changes over the growing season, and any noteworthy differences in the rate of change at different times. Thereafter, the magnitude and effect size of defoliation responses are compared across variables to assess the relative sensitivity of different growth components.

### Above-ground biomass

In control plants, above-ground dry matter (AGDM) increased rapidly from early summer (mid-November), peaked in early February, and then declined steadily as growth senesced to reach its lowest level in late winter (July) (Figure 1). In contrast, leaf growth in defoliated plants remained strongly suppressed, showing only a slight increase towards the end of the growing season before gradually declining back to near its initial level by late July. During the critical late-summer to autumn period, when geophytes typically replenish their USO carbohydrate reserves, the maximum growth rate of control plants was nearly 23-fold higher than that of defoliated plants, with defoliated plants attaining 84.1% less AGDM than controls at their peaks in early February (Table 1).

**Figure 1.**
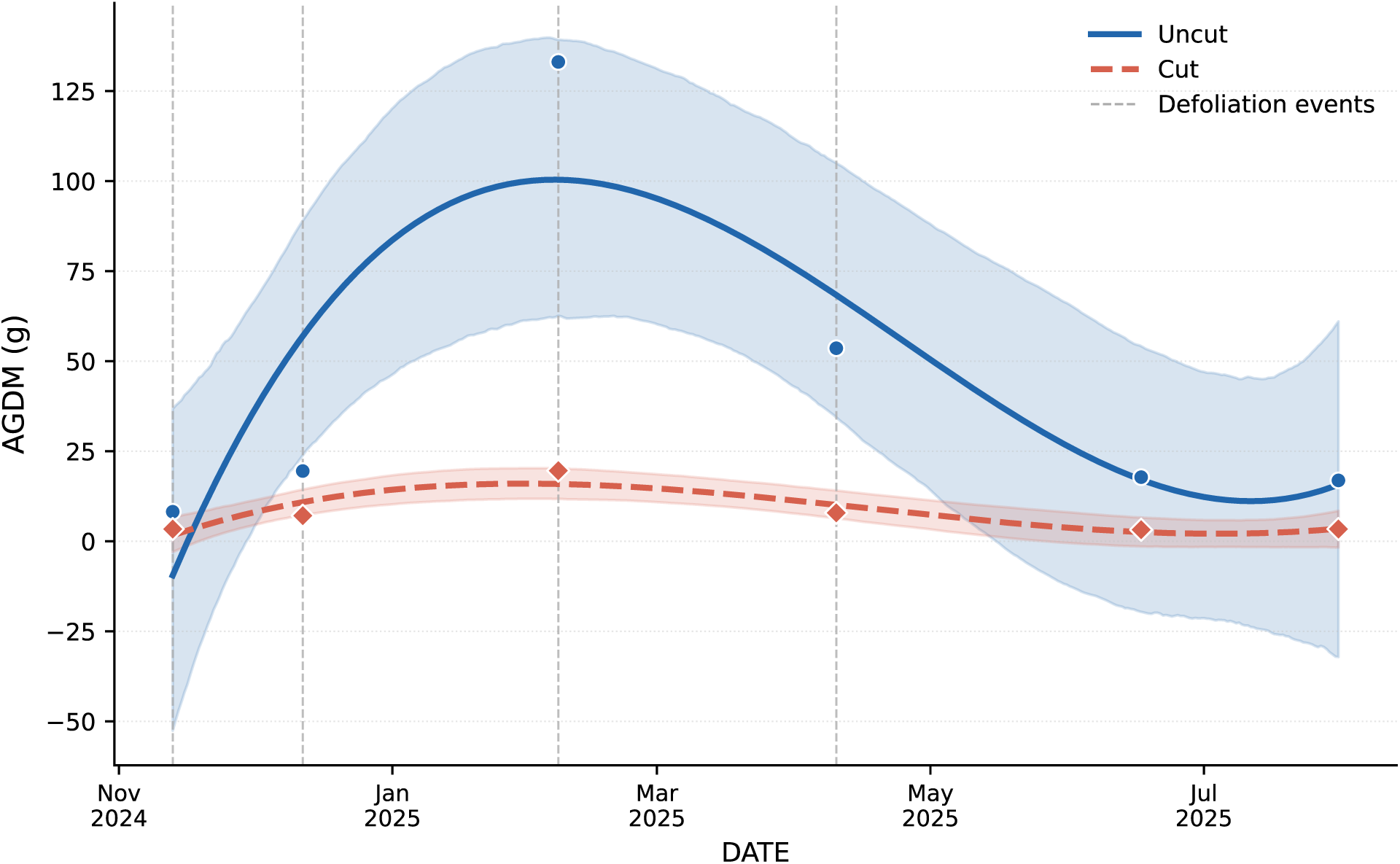
Seasonal dynamics of leaf above-ground dry matter (AGDM) of *Eucomis autumnalis* in defoliated and undefoliated control plants, modelled using Generalised Additive Models (GAMs) with 95% bootstrap confidence intervals. Vertical dashed grey lines indicate the four defoliation events.

**Table 1.** Maximum modelled rates of change, peak values, and relative differences for all measured above-ground and bulb variables of *Eucomis autumnalis* during the critical window for bulb biomass and starch replenishment (February–May 2025). AGDM = above-ground dry matter.

| Variable | UNCUT max rate (date) | CUT max rate (date) | Ratio | UNCUT peak value | CUT peak value | % difference |
| --- | --- | --- | --- | --- | --- | --- |
| **AGDM (g)** | +0.529 g day<sup>-1</sup> (1 Feb) | +0.023 g day<sup>-1</sup> (1 Feb) | Uncut 22.8× faster | 100.4 g (6 Feb) | 16.0 g (1 Feb) | –84.1% |
| **Bulb mass (g)** | +0.786 g day<sup>-1</sup> (18 Feb) | +0.389 g day<sup>-1</sup> (21 Mar) | Uncut 2.0× faster | 214.7 g (30 May) | 95.2 g (30 May) | –55.7% |
| **Starch content (%)** | +0.154 % day<sup>-1</sup> (19 Apr) | +0.284 % day<sup>-1</sup> (30 May) | Cut 1.8× faster | 35.2% (30 May) | 28.0% (30 May) | –20.4% |
| **Starch pool (g)** | +0.421 g day<sup>-1</sup> (25 Mar) | +0.233 g day<sup>-1</sup> (30 May) | Uncut 1.8× faster | 76.8 g (30 May) | 25.6 g (30 May) | –66.7% |
| **Water content (%)** | –0.044 % day<sup>-1</sup> (30 May) | +0.085 % day<sup>-1</sup> (1 Feb) | Opposite directions | 38.0% (1 Feb) | 48.8% (4 Feb) | +28.4% |
| **Water pool (g)** | +0.102 g day<sup>-1</sup> (1 Feb) | –0.169 g day<sup>-1</sup> (30 May) | Opposite directions | 58.5 (30 May) | 42.0 g (1 Feb) | –30.0% |

### Bulb biomass

Bulb dry mass initially decreased over the first winter (June–September 2024), then, in control plants, followed a clear hump-shaped seasonal trajectory, increasing steadily through the growing season to a peak in July 2025 before gradually decreasing through to October 2025 (Figure 2a). In sharp contrast, bulb biomass in defoliated plants dipped after the second defoliation, recovering somewhat in late autumn to reach a winter peak in 2025, thereafter declining gradually through to the end of the experimental period in October 2025, culminating in a final biomass deficit of approximately 130 g relative to control bulbs (Figure 2a). During late summer to autumn, bulb growth in defoliated plants was more than twice as slow as in controls, with peak mass 55% lower than that of control plants over the same period (Table 1).

**Figure 2.**
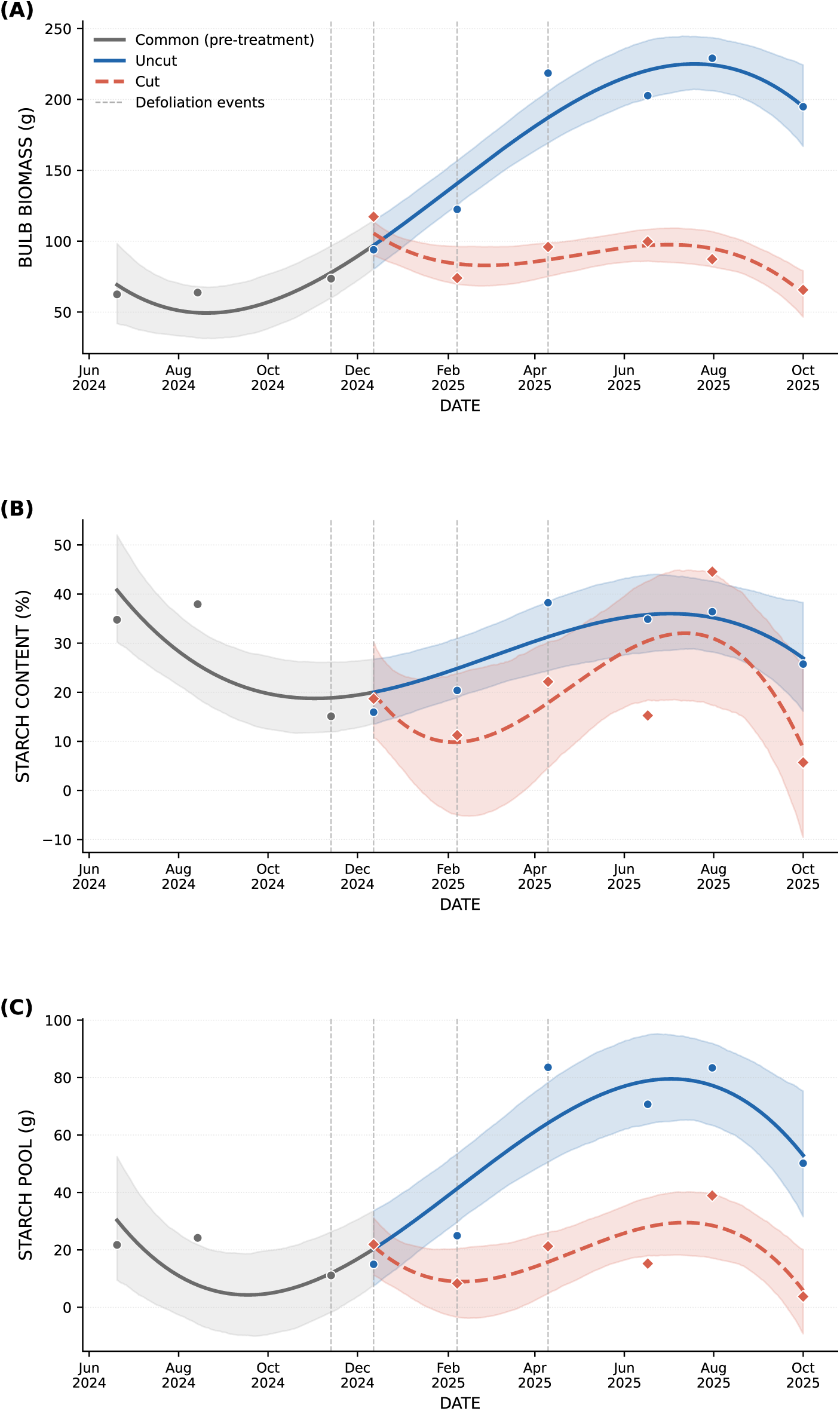
Seasonal dynamics of (A) bulb dry mass, (B) starch concentration, and (C) starch pool of *Eucomis autumnalis* in defoliated and undefoliated control plants, modelled using Generalised Additive Models (GAMs) with 95% bootstrap confidence intervals. A single common smooth (grey) was fitted through the shared pre-treatment observations.

### Bulb starch content

Bulb starch concentration declined from just over 40% at the start of the experiment in June 2024 to approximately 20% by the end of spring in October (Figure 2b). Thereafter, starch levels recovered steadily through the growing season and autumn, peaking at over 35% in mid-winter 2025, before declining again in spring 2025. In defoliated plants, starch concentration dropped sharply after the second defoliation to just above 10%, but recovered 1.8 times faster than in control plants during autumn (Table 1), reaching a lower mid-winter peak and showing a similar decline the following spring. Starch concentration showed substantial variability in both treatments, particularly in defoliated plants, with overlapping confidence intervals throughout the post-defoliation period (Figure 2b), suggesting that the observed differences should be interpreted with caution rather than as statistically distinct seasonal trends.

### Bulb starch pool

Seasonal changes in total bulb starch pool in both control and defoliated plants, calculated from bulb mass and starch concentration, closely mirrored changes in bulb mass (Figure 2c) rather than starch concentration, as bulb mass trajectories diverged strongly between treatments while starch concentration remained comparatively similar. By the end of May, when both above- and below-ground growth had tailed off, defoliated plants had a total starch pool two-thirds smaller than control plants (Table 1).

### Bulb water content

Bulb water concentration declined from winter 2024 through spring into early summer. In control plants, this decline continued through summer and autumn, reaching a minimum of 26% in winter 2025 before recovering to 40% in spring (Figure 3a). Notably, defoliated plants initially showed the opposite pattern during the defoliation period, with bulb water concentration rising rapidly to nearly 50% in early February, compared with 38% in control bulbs (Table 1), before declining to just under 35% by winter 2025 and rising again to 55% in spring. Defoliated plants consistently had higher bulb water concentrations than controls from summer through spring.

**Figure 3.**
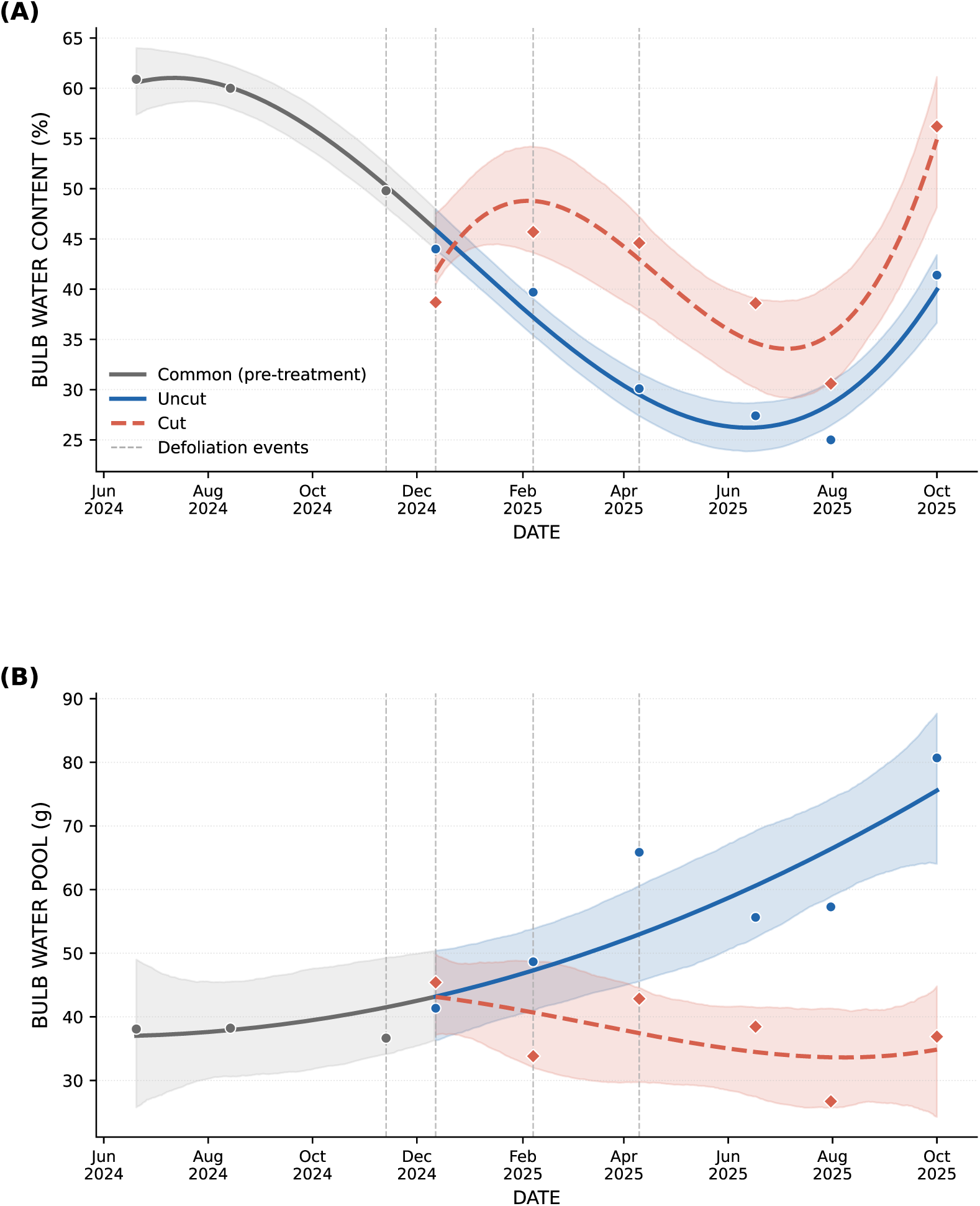
Seasonal dynamics of (A) bulb water content and (B) bulb water pool (g) of *Eucomis autumnalis* in defoliated and undefoliated control plants, modelled using Generalised Additive Models (GAMs) with 95% bootstrap confidence intervals. A single common smooth (grey) was fitted through the shared pre-treatment observations.

### Bulb water pool

Total bulb water pool in both control and defoliated plants tracked seasonal changes in bulb mass more closely than changes in water concentration (Figure 3b). Control plants showed a steady increase throughout the experimental period, whereas defoliated plants declined after defoliation to the end of the experiment in October 2025. During autumn and across the experimental period, defoliated bulbs had total water pools that were 30% and 35.1% lower, respectively, than those of control plants (Tables 1 and 2).

**Table 2.** Time-integrated area under the GAM-fitted curve (AUC) for all variables measured on *Eucomis autumnalis* over the period during which the effects of defoliation (CUT) versus undefoliated plants (UNCUT) were measured. AUC units are the variable unit × day. AGDM = above-ground dry matter.

| Variable | UNCUT AUC | CUT AUC | % difference |
| --- | --- | --- | --- |
| AGDM (g·day) | 14,549 | 2,372 | –83.7% |
| Bulb mass (g·day) | 54,073 | 26,123 | –51.7% |
| Starch content (%·day) | 8,872 | 6,003 | –32.3% |
| Starch pool (g·day) | 17,650 | 5,485 | –68.9% |
| Bulb water content (%·day) | 9,533 | 12,337 | +29.4% |
| Bulb water pool (g·day) | 16,713 | 10,851 | –35.1% |

### Relative defoliation responses and effect sizes

Defoliation most strongly reduced above-ground production, with seasonal integrated AGDM 83.7% lower than controls, compared with a 51.7% reduction in bulb mass (Table 2). Among the bulb content variables, starch pool was depleted the most by defoliation, declining by 68.9%, followed by bulb mass (−51.7%) and bulb water pool (−35.1%) (Table 2). Starch concentration was less responsive (−32.3%), while bulb water concentration increased by 29.4% under defoliation.

Effect size analysis indicated that below-ground reserves were the most responsive to defoliation relative to variability, with bulb mass, starch pool, and bulb water pool all showing very large treatment effects (d = 1.298–1.381; Table 3), indicating high sensitivity to defoliation. Defoliation had a large effect on AGDM (d = 0.823). In contrast, its effects on starch concentration were small (h = 0.235), while bulb water concentration showed a negligible inverse effect (h = −0.163; Table 3)

**Table 3:** Effect sizes measured by Cohen’s h for proportion variables (starch content %, water content %) and Cohen’s d for mass and pool quantities of the bulbs of *Eucomis autumnalis*.

| Variable | Metric | UNCUT mean | CUT mean | Effect size | Magnitude | Direction |
| --- | --- | --- | --- | --- | --- | --- |
| AGDM (g) | Cohen's *d* | 41.5 g | 7.4 g | +0.823 | Large | Uncut > Cut |
| Bulb mass (g) | Cohen's *d* | 177.0 g | 90.0 g | +1.376 | Very large | Uncut > Cut |
| Starch pool (g) | Cohen's *d* | 54.6 g | 18.2 g | +1.381 | Very large | Uncut > Cut |
| Bulb water pool (g) | Cohen's *d* | 58.2 g | 37.4 g | +1.298 | Very large | Uncut > Cut |
| Starch content (%) | Cohen's *h* | 28.6% | 19.6% | +0.235 | Small | Uncut > Cut |
| Bulb water content (%) | Cohen's *h* | 34.6% | 42.4% | –0.163 | Negligible | Cut > Uncut |

*Benchmarks: |d| or |h| < 0.2 negligible, 0.2–0.5 small, 0.5–0.8 medium, 0.8–1.2 large, > 1.2 very large.*

## DISCUSSION

Understanding how grassland geophytes acquire, store, and remobilise resources such as carbon, minerals, and water in response to seasonal variation and recurrent disturbances (e.g. frost, fire, drought, and herbivory) is crucial for developing management strategies that support their recovery and persistence. While geophyte seasonality has been studied in the winter-rainfall (Cape) region of South Africa, where plants endure long, hot, dry summers (Ruiters and McKenzie 1994; Ruiters 1995; Daniels et al. 2014), and in temperate and Mediterranean grasslands elsewhere, where severe winter cold and drought are major constraints (e.g. Al-Tardeh et al. 2008; Janeček et al. 2015; Shane and Pate 2015; Bernard et al. 2020), the seasonal dynamics of underground storage organs (USOs) in the mesic subtropical and temperate grasslands of southern Africa remain largely unknown.

Nearly a century ago, Bews and Vanderplank (1930) reported relatively stable carbohydrate and water concentrations throughout the year in corms of *Hypoxis hemerocallidea* dug up from a grassland near our study site, with only modest reserve accumulation before dormancy, slight depletion during spring regrowth, and a small increase in water content in spring. The only other seasonal data for mesic South African mesic grassland geophytes come from *Merwilla plumbea* sampled on four occasions, where starch and non-structural carbohydrate reserves declined sharply in spring before recovering strongly during summer (Morris and Nkuna 2025b). Our study provides the first detailed seasonal account of carbohydrate and water dynamics in a mesic grassland geophyte, identifying periods when the bulb functions alternately as a source of energy for growth and a sink for reserve accumulation before overwintering.

Our findings help address a substantial knowledge gap regarding the impacts of herbivory on African grassland forbs, which can be severe and long-lasting (Scott-Shaw and Morris 2015; Veblen et al. 2016; Chamane et al. 2017a, 2017b; Morris and Scott-Shaw 2019). Although previous studies have documented declines in forb populations and diversity under chronic grazing, the mechanisms by which grazing and trampling reduce forb vigour and persistence remain unclear. Morris (2021) proposed that repeated removal of photosynthetic tissue reduces spring growth and USO size, progressively imperiling plants as these effects accumulate (Morris and Nkuna 2025a). Our study adds an important mechanistic detail: growing-season defoliation impairs energy storage not by reducing the bulb’s capacity to concentrate starch but by preventing bulbs from attaining sufficient size to accumulate meaningful absolute reserves. Thus, it is the size of the storage base, rather than storage efficiency per unit tissue, that collapses under defoliation. The resulting stunted bulbs likely hold too little energy reserve to support further spring or summer regrowth under persistent grazing, consistent with the “bulb bankruptcy” model we propose.

Below, we first discuss how late-season defoliation disrupts the critical period of starch accumulation, then how a shift from sink to source relationships in bulbs leads to structural loss of storage capacity and signs of impending “bulb bankruptcy”, and finally the implications of these dynamics for the management and persistence of grassland geophytes under chronic grazing.

### Late-season storage disruption

Morris and Nkuna (2025b) showed that neither single spring nor double spring–summer defoliation reduced aboveground growth. Plants compensated for tissue loss by mobilising bulb starch reserves, enabling rapid leaf regrowth and maintaining aboveground biomass, bulb mass, and NSC and starch levels comparable to undefoliated plants by the end of the growing season. Based on these seasonal dynamics, they predicted that grazing during the critical late-summer to autumn replenishment period, when growth slows, and plants mature and senesce, would have much stronger negative effects by disrupting the replenishment of bulb carbohydrate reserves required for future growth and persistence (Morris and Nkuna 2025b).

Our results support this prediction, showing that the late-summer to autumn period is critical for resource storage and sustained plant growth. Bulbs of undefoliated plants accumulated biomass and replenished starch most rapidly during this period, reaching peak levels in winter. In contrast, aboveground growth peaked at the end of the summer growing season (February) and declined thereafter as growth slowed and leaves eventually senesced. These patterns indicate a shift in allocation toward below-ground storage during autumn.

This pronounced shift from above-to below-ground growth and storage during the critical autumn period, when above-ground biomass declines and leaf senescence begins, is consistent with seasonal dynamics of geophyte growth and energetics observed in other studies. A global synthesis by Martínez-Vilalta et al. (2016) and a study of 80 temperate meadow species by Lubbe et al. (2021) confirm that this late-season investment in belowground reserves is a widespread pattern, with carbohydrate concentrations in underground storage organs (USOs) typically reaching their lowest levels during active vegetative growth (e.g. Mzabri et al. 2025) and peaking during autumn and winter as growth slows and dormancy approaches. Along this phenological trajectory, Meyer and Hellwig (1997) showed that maximum rhizome starch levels in geophytes occur immediately before leaf yellowing, while Marvel (1986) reported a near-doubling of storage reserves just before foliar senescence. In economic terms, this phase represents the critical consolidation of capital (Chapin et al. 1990), when surplus carbon is converted into underground structural stores. Khodorova and Boitel-Conti (2013) provide a mechanistic explanation for these winter peaks, demonstrating that physiological processes within bulbs actively promote reserve accumulation and organ enlargement in preparation for future growth. Generally, temperature rather than photoperiod controls seasonal growth, storage, dormancy, and flowering cycles in geophytes (Kumari et al. 2024).

The shape of seasonal reserve depletion and replenishment patterns varies among species, reflecting different storage strategies and contributing to differences in tolerance to defoliation and other disturbances (Menke and Trlica 1981). Morris and Nkuna (2025b) found that spring and summer defoliation caused only a temporary change in the shape of seasonal energy storage patterns in a mesic grassland geophyte. In contrast, our results show that when repeated defoliation continued into autumn, starch accumulation per bulb cell still increased, but overall bulb growth was reduced by more than half during this period, leading to a marked suppression of total starch reserve accumulation. This coincided with a more than 80% reduction in AGDM relative to controls, substantially limiting leaf area available for carbon capture during the critical storage period. During this period, low temperatures restrict regrowth, but photosynthesis can still occur, making late-season leaf area particularly important for reserve accumulation (Anderson and McNaughton 1973; Körner 2015).

Few studies have been undertaken on the effect of defoliation, especially during autumn, on carbohydrate storage in grassland geophytes (Morris 2021; Morris and Nkuna 2025a). Wang et al. (2016) similarly showed that autumn defoliation reduced non-structural carbohydrate reserves in storage crowns and roots, with subsequent reductions in spring regrowth. Likewise, in an herbaceous shrub (lucerne), repeated defoliation, including autumn, significantly reduced soluble sugar and starch reserves in taproots and crowns, resulting in lower perennial biomass and reduced productivity (Teixeira 2006). Together, our results and these findings highlight the sensitivity of belowground carbon storage to disturbance during late-season reserve accumulation, underscoring the need for more experimental work focused specifically on defoliation effects within the critical autumn storage period and a better understanding of the role of this critical autumn storage period in maintaining the source-sink energy dynamics of the whole-plant.

### Seasonal source-sink dynamics

Our study characterised the seasonal source–sink dynamics of *E. autumnalis* and showed how these are fundamentally reorganised by repeated defoliation. Three distinct phases were evident in the undisturbed controls. First, during spring mobilisation, starch stored in the bulb served as the principal energy source for early vegetative growth, evident by declining bulb biomass and starch pools while new leaves expanded. Second, during maximum vegetative growth, growth was likely sustained primarily by concurrent photosynthesis in the expanding leaf canopy. During this period, below-ground replenishment lagged behind above-ground development: above-ground dry matter peaked in late summer, whereas bulb biomass and total starch pools continued to increase. Third, a late-season sink-loading phase developed as vegetative growth slowed and the canopy senesced, with photosynthates increasingly redirected below ground. This resulted in maximum bulb size and starch reserves by mid-winter before these reserves were mobilised again to support the following spring flush.

Such pronounced seasonal oscillations in source–sink relationships are characteristic of plants in strongly seasonal environments, where opportunities for photosynthesis and growth vary markedly through the year (Mooney 1972; Martínez-Vilalta et al. 2016). Underground storage organs (USOs) play a pivotal role in this cycle by supplying carbohydrates, including starch hydrolysed to soluble sugars (Franková et al. 2003; Raheem and Thoss 2016), which are translocated to actively growing meristems (Lapointe 2001) to support the strong sinks created by emerging shoots, leaves, flowers and roots (e.g. Ruiters and McKenzie 1994; Naor et al. 2008; Yoshida 2021; Morris and Nkuna 2025b). Experimental evidence from a southern African mesic grassland geophyte likewise demonstrates the dependence of spring growth on stored reserves (Morris 2021). As the season progresses and above-ground growth slows, USOs transition from carbon sources to carbon sinks, accumulating carbohydrates and mineral reserves for the following growing season, as reflected in the increasing bulb starch pools observed here and documented widely elsewhere (e.g. Orthen and Wehrmeyer 2004; Martínez-Vilalta et al. 2016; Petrussa et al. 2018; Pouris et al. 2020; Pouris et al. 2022).

Recurrent defoliation reversed the seasonal source–sink pattern, forcing the bulb to repeatedly supply energy for regrowth rather than store carbohydrates. As a result, starch concentrations in bulb tissues declined during summer and only recovered in autumn, while bulb growth and total underground starch pools remained suppressed. Carbon was continuously diverted to rebuilding lost photosynthetic tissue, so regrowth became the dominant sink, effectively preventing reserve accumulation.

Compared with grasses, relatively few studies have examined how above-ground defoliation alters below-ground growth and reserve dynamics in grassland forbs, including geophytes. Our previous work on mesic grassland forbs shows that repeated summer defoliation substantially depletes underground storage organs, with effects compounding across seasons (Morris and Nkuna 2025a). Likewise, rhizomes of temperate geophytes are disproportionately reduced relative to shoots under repeated summer cutting (Ottaviani et al. 2021). In contrast, some studies report that defoliation can, under certain regimes, promote longer-term increases in storage allocation through compensatory regrowth and reserve rebuilding (Janeček and Klimešová 2014), alongside shifts toward more acquisitive root and leaf traits that may enhance tolerance to future disturbance (Martínková et al. 2026). However, other evidence shows that shoot removal can rapidly deplete below-ground biomass and starch while altering the carbon–nitrogen balance in storage organs, reflecting mobilisation of reserves to sustain regrowth (Kleijn et al. 2005). Together, these findings highlight complex and often contrasting subterranean responses to defoliation in forbs (Siebert et al. 2024), with repeated or chronic biomass loss potentially driving progressive depletion of storage organs and raising the risk of eventual “bulb energy bankruptcy” under sustained grazing pressure.

### Signs of bulb bankruptcy

Repeated defoliation of *E. autumnalis* produced a suite of symptoms analogous to those of a business approaching insolvency (Chapin et al. 1990). In an intact plant, stored reserves function as capital stock: starch accumulated in the bulb finances spring growth, while leaves generate photosynthetic income that repays this capital and supports new storage. Any surplus is reinvested as capital accumulation in the bulb before dormancy. This reserve system operates within a coordinated above- and belowground economic framework, integrating water-use traits and storage organs into a whole-plant strategy (Lubbe et al. 2021). Under repeated defoliation, this system progressively failed. Income flow declined as leaf area was repeatedly removed, while expenditure on canopy replacement increased after each defoliation. At the same time, the capacity for capital accumulation was reduced, resulting in a persistent failure to rebuild underground reserves.

Several indicators point to this impending state of “bulb bankruptcy” in defoliated *E*. *autumnalis*. The most obvious was the collapse in the bulb’s capital stock: bulb biomass and starch pools were markedly diminished, despite only modest reductions in starch concentration within individual storage cells. Thus, rather than a loss of physiological storage function, the system failed through a reduction in storage capacity itself. The storage account remained operational, but too small to sustain additional growth. Plant growth may even cease before reserves are fully exhausted (Bloom et al. 1985).

A second, and somewhat unexpected, symptom was the contrasting behaviour of bulb water content. Underground organs in savanna forbs can function as significant water reservoirs supporting persistence under drought (da Silva and Rossatto 2019). In our study, defoliated bulbs consistently contained a higher proportion of water. This likely reflects continued water uptake despite reduced transpirational demand following repeated canopy removal during the growing season. However, because bulb size—and thus total storage capacity—was markedly reduced, the absolute stored water capital was also diminished. The result was small, water-rich bulbs rather than large, dense, starch-rich storage organs characteristic of intact plants. In financial terms, the system appears to have entered a form of “liquidation”, in which remaining capital was redistributed into low-return liquid reserves that could not sustain future growth. This decoupling between structural attributes and functional storage capacity is consistent with evidence that dry matter-based traits may not reliably track carbon investment or allocation across plant organs (Ottaviani et al. 2026).

Ultimately, bulb bankruptcy reflects not only depletion of carbohydrate reserves but a progressive collapse of the entire source–sink economy. Once the leaf canopy can no longer generate sufficient income to replenish underground capital stock, the system enters a deficit state in which maintenance and replacement costs exceed returns. Chronic defoliation thus drives geophytes into an energetic downward spiral from which recovery becomes increasingly difficult if herbivory is sustained. This collapse may also involve active structural reorganisation and senescence of belowground storage tissues, including programmed degeneration and functional abandonment of ageing clonal organs (da Silva et al. 2026), suggesting that “loss of capital” reflects both depletion and regulated divestment from unproductive assets.

Recognising these early signs of declining capital efficiency in geophytes has important implications for tailoring grazing management to sustain diverse populations of mesic grassland forbs.

### Implications for grassland management

Wang et al. (2016) emphasised that maintaining a full leaf canopy during the growing season is critical for replenishing the carbohydrate reserves stored in crowns required to ensure winter survival and spring growth. However, when grasslands are grazed, especially by concentrated herds of livestock (Franke and Kotzé 2022), few forbs (<10%), including even rare species, escape defoliation or mechanical leaf damage from the hooves of livestock (Chamane et al. 2017a). While reducing overall stocking rates and densities can lessen defoliation pressure and the risk to forbs, recovery ultimately depends on providing adequate rest periods that allow plants to rebuild reserves in USOs. In particular, our results clearly indicate that seasonal resting during the late-season replenishment phase is essential for geophytic forbs. Periodic (rotational) full-year rests, encompassing the growing season and autumn (McDonald et al. 2019), are already recommended to maintain the vigour of grazed grasses (Kirkman et al. 2023, Fynn et al. 2026). Such long rests are likely to also benefit forb populations by allowing uninterrupted completion of the seasonal source–sink cycle, but this hypothesis requires further testing under both field and controlled conditions.

## CONCLUSION

This study provides the first detailed mechanistic evidence of how repeated defoliation disrupts the seasonal carbon economy of a mesic grassland geophyte. Rather than reducing the bulb’s physiological capacity to accumulate starch, repeated defoliation prevented bulbs from growing large enough to build adequate statch reserves. By repeatedly diverting carbon from storage to leaf replacement, defoliation reversed the normal seasonal source-sink dynamics, progressively reducing bulb biomass, starch pools, and water reserves. These responses provide strong support for the concept of “bulb bankruptcy”, whereby repeated canopy loss progressively erodes the capital required to sustain future growth.

These findings identify late summer and autumn as the critical period for replenishing underground reserves. Because grassland forbs cannot escape repeated defoliation and trampling wherever livestock are present, management should focus on reducing the frequency and severity of damage through moderate stocking densities and periodic rotational recovery rests. Providing uninterrupted rest during the autumn reserve-replenishment period, together with occasional full-season rests, is likely to be essential for maintaining the vigour and persistence of *Eucomis autumnalis* and, potentially, many other geophytes in grazed mesic grassland.

## ACKNOWLEDGEMENTS

We are grateful to Welcome Ngcobo for his help in setting up and maintaining the experiment, and Anita Morris for her support with treatment application and data collection

